# Chromatin phase separated nanoregions regulated by cross-linkers and explored by single particle trajectories

**DOI:** 10.1101/2022.12.16.520776

**Authors:** A Papale, D. Holcman

## Abstract

Phase separated domains (PSDs) are ubiquitous in cell biology, representing nanoregions of high molecular concentration. PSDs appear at diverse cellular domains, such as neuronal synapses but also in eukaryotic cell nucleus, limiting the access of transcription factors and thus preventing gene expression. We study here the properties of PSDs and in particular how they can be generated by polymers. We show that increasing the number of cross-linkers generate a polymer condensation, preventing the access of diffusing molecules. To investigate how the PSDs restrict the motion of diffusing molecules, we estimate the mean residence and first escaping times. Finally, by computing the mean square displacement of single particle trajectories, we can reconstruct the properties of PSDs in term of a continuum range of anomalous exponents. To conclude, PSDs can result from a condensed chromatin, where the number of cross-linkers control the molecular access.

---

Chromatin in the cell nucleus is organized uniformly (euchromatin), forming regions associated with gene expression, or in dense heterogeneous regions called heterochromatin, where genes are hardly expressed [1]. Heterochromatin is less accessible to transcription factors [2], remodelers or polymerase. However, the formation and maintenance of heterochromatin microdomains remain unclear, although remodelers such as histone HP1, NURD remodelers or transcription factors can bind chromatin to form local foci through specific interactions [3–5] and can also modify the local condensation. Foci can also be generated during double-stranded DNA break [6, 7], the property of which can be revealed by single particle trajectories (SPTs). In the case of tagged NURD remodeler, SPTs reveal chromatin organization, where decondensation is associated with an increase of the anomalous exponent [8–10], a parameter that quantifies how the mean square displacement depend on the time increment. This decondensation is associated to increase of the confinement length, that characterizes the confined volume (in 3d) or the surface (in 2d) visited by trajectories.

Phase Separated Domains (PSDs) [11–13] are regions with a size of hundreds nanometers, that can found in cell biology ranging from neuronal organizations [14], post-synaptic density, synaptic organization [14, 15], immune synapses or nucleus organization, possibly originated from disorder aggregates [16–18], or local chromatin interaction [19–22]. Motions in PSDs is often characterized by a large range of transient to permanent trappings, that can be characterized by potential wells [18]. Chromatin is also organized in large dense regions called Topological Associated Domains (TADs), regions with enhanced local interactions, revealed by population analysis of Hi-C maps at Mbps scale. It remains unclear how PSDs affect the dynamics of stochastic particles and how the exchange rate is controlled across.

We explore here how PSDs can be generated and regulate the the in and outflux of diffusing molecules. Several polymer models have been used to investigate the spatial organization of chromatin [23] at various scales, including TADs, based on diffusive binders with specific binding sites [24–26], attractive interactions among epigenomic domains [22, 27, 28] or random cross-linkers [29–31]. Using cross-link polymer models, we explore how local high density chromatin regions can emerge and form PSD. To quantify the ability to prevent molecular exchange, we explore how diffusing molecules can be excluded from PSDs due to spatial constraint and volume exclusion. By increasing the number of cross-linkers, PSDs emerge and the reduced volume inside the condensed chromatin can prevent most diffusing molecules from accessing. We characterize the PSDs by estimating a penetration length across their fuzzy boundary. To quantify the porosity of the PSD boundary to Brownian molecules, we compute the mean residence time and the first escaping times [32, 33]. The deviation from diffusion due to chromatin organization is revealed by the spectrum of anomalous exponent computed over SPTs, that decays from the center to the periphery and also by increasing the number of connectors.

## Modeling chromatin phase separation with a Random-cross-link Polymer model

To investigate how chromatin condensation can generate phase-separated domains, we generalize the cross-linker polymer model (RCL) [29, 34, 35], which consists of the Rouse model with randomly added cross-linkers but fixed for a given configuration. We adopted a coarsegrained semi-flexible chain with volume-excluded interactions modeled by Lennard-Jones forces, following the Kremer-Grest bead-spring polymer model [36, 37]). Each *N_mon_* – monomer represents a segment of 3 kbps with a size of *σ* = 30*nm*, and additional cross-linkers are chosen at random positions as in the RCL-polymer model [29, 34] (Supp. Material). A cross-linker consists of a harmonic spring between two randomly chosen monomers (fig. 1A). The chromatin network resulting from *N_c_* random connectors defines a realization and accounts for the local organization induced by cohesin, condensin or CTCF and thereby combination [30, 38–40].

**FIG. 1.**
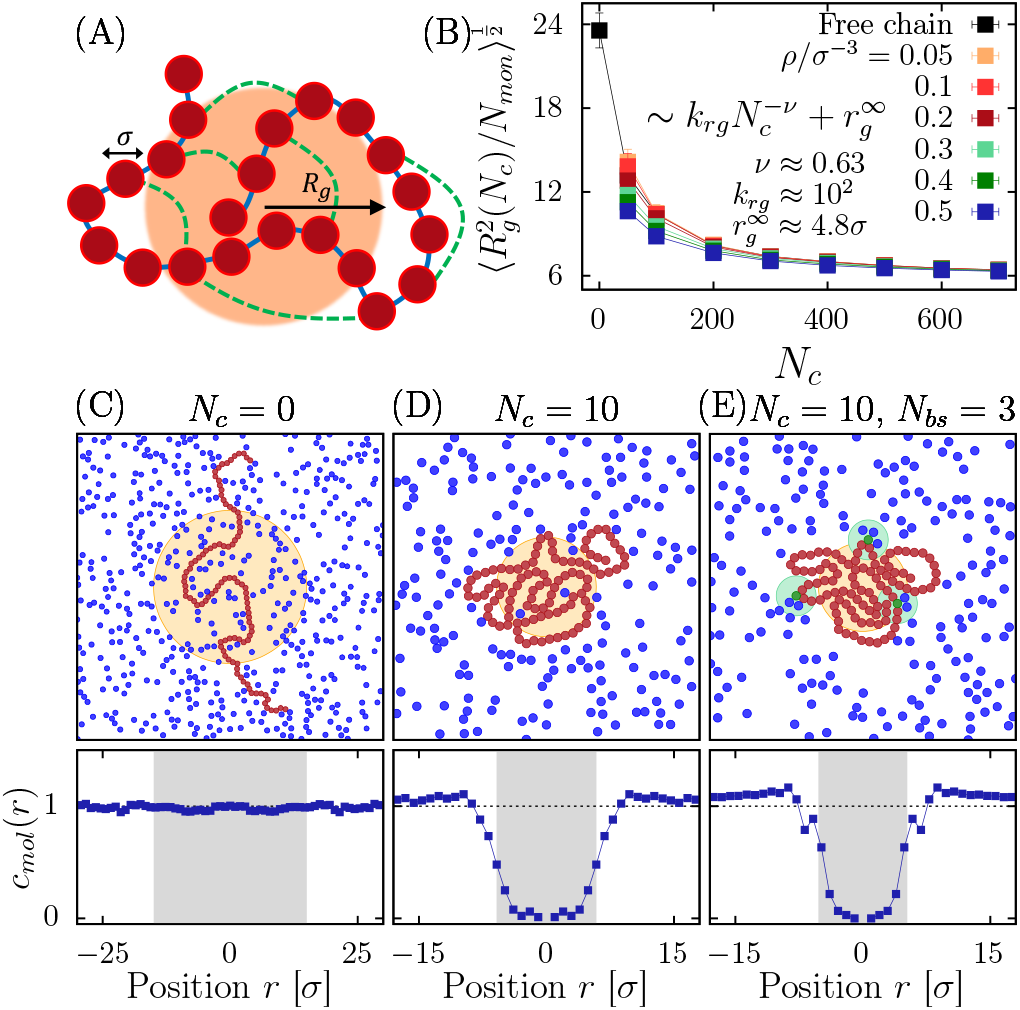
**A.** Scheme of local chromatin reconstruction based on a cross-linked polymer model (red bead of diameter σ) connected by springs (blue) with random connectors (green dots). The ball *B*(*R_g_*) (orange domain) defines the radius gyration. **B.** Mean gyration radius vs number of random cross-linkers *N_c_*, for various densities *ρ*: a smooth transition occurs from a swollen chain to a compact state (*N_mon_* = 2000). **C-E.** Linear chain (red monomers) without random connectors embedded in *N_mol_* Brownian molecules (blue). Random connectors drive the free particles outside *B*(*R_g_*). When there are *N_bs_* binding sites, the concentration of molecules *c_mol_*(*r*) at distance *r* from the center, is depleted in *B*(*R_g_*) (lower panels).

We first investigate the effects of increasing the number of random cross-linkers on an isolated chain revealing a transition from a coil configuration to a globular state, as characterized by the gyration radius 〈*R_g_*〉 (fig. 1B, black curve) [34, 41] where 〈.〉 represents the average over simulations and cross-linkers realizations. We found that gyration radius is well approximated by a power-law 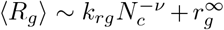, where *k_rg_* = 10^2^ ± 20 *σ*, *ν* = 0.63 ± 0.05, 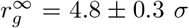. In the limit of large amount of connectors, *N_c_* → ∞, 〈*R_g_*〉 converges to a non-zero constant value 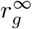 due to the volume-excluding interactions.

To investigate how the chromatin structure can influence the dynamics and the distribution of random moving molecules, we simulated a RCL-chain with *N_mon_* = 2000 monomers and *N_c_* = {50, 100,.., 700} random connectors embedded in a volume containing *N_mol_* = 8000 diffusing molecules that interact with the chromatin via Lennard-Jones volume exclusion forces (Supp. Material), fig. 1CD.

We also introduce specific attractive interactions between diffusing molecules and a set of *N_bs_* = {0, 10} selected monomers of the chain (fig. 1E). We performed fixed-volume molecular dynamics simulations [42] (Supp. Material) in a fixed cubic volume *V* with periodic boundary conditions and the overall density is defined by *ρ* = (*N_mon_* + *N_mol_*)/*V* and *ρ*/*σ*^3^ ∈ [0.05, 0.5].

We report that the average gyration radius 〈*R_g_*〉 is slightly affected by the presence of the diffusing molecules (fig. 1B), in particular for small *N_c_*, the effective density of the polymer increases. The nano-region occupied by the polymer varies dynamically with the chain motion thus we define the boundary of the separated phase domain as the convex ball Ω = *conv*({**r**|**r** – **r**_CM_| ≤ 〈**R_g_**〉, }, where **r**_CM_ is the polymer center of mass and the radius is 〈*R_g_*〉. Interestingly, diffusing particles can be excluded from the region Ω as the number of cross-linkers is increasing (fig. 1C-D below) even with binding domains (fig. 1E below).

## Statistics distribution of diffusive molecules with in a PSD

To study the distribution of Brownian molecules with respect to the PSD, we use as a reference the radial distribution of molecules with respect to the center of mass *CM*

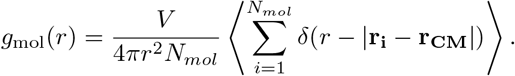

Similarly, the distribution of monomers is characterized by

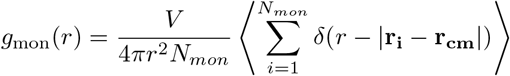

and the pair correlation function molecules-monomers is given by

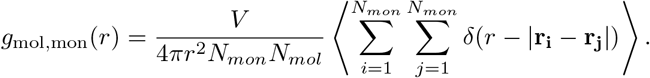

The radial distribution functions of monomers and molecules reveal that the RCL-chain separates diffusing molecules, a phenomena that is amplified by increasing the number of random connectors (fig. 2A-B), regardless of the overall density (see also SI Fig. S1 for the radial pair distribution functions for various density *ρ*). We thus conclude that the presence of random connectors can create a separation between a condensed polymer and interacting molecules.

**FIG. 2.**
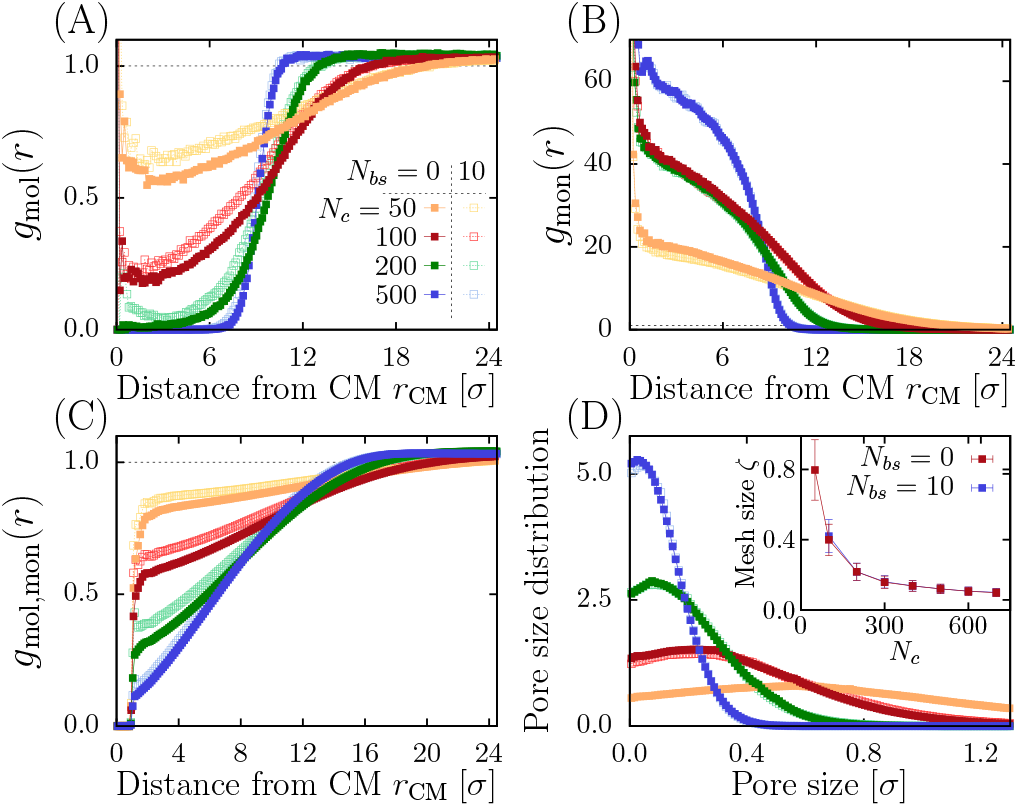
**A.** Molecular radial distribution function *g*_mol_(*r*) for various *N_c_* ∈ {50, 100, 200, 500} at density *ρ* = 0.05*σ*^-3^, compared to the refence constant dashed line. Full (resp. empty) symbols indicate cases with *N_bs_* = 0 (*N_bs_* = 10). **B.** Polymer radial distribution function *g*_mon_(*r*). **C.** Molecules-monomers pair correlation function *g*_mol_(*r*). **D.** Pore size distribution. Inset: average mesh size *ζ* vs. *N_c_*.

To further characterize the spatial organization of the RCL-chain, we analyze the available space for diffusion in the region Ω [43, 44] by estimating the pore size distribution *P_s_* from the maximum volume that do not contain any other monomer inside the region (fig. 2D). The mesh size is defined as the mean pore radius *ζ* = 〈*s*〉 = ∫ *sP_s_ds* that can be approximated as 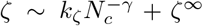. For *N_bs_* = 0 (resp. *N_bs_* = 10) fitting the simulations reveals an exponent *γ* = 1.13 ± 0.01 (1.23 ± 0.04), *k_ζ_* = 60 ± 3 *σ* (100 ± 20 *σ*) and ζ^∞^ = 0.062 ± 0.002 *σ* (0.071 ± 0.004 *σ*) (fig. 2D inset). To conclude, increasing the connectors *N_c_* forces the polymer to condense and to progressively exclude random particles, sharpening the boundary of the PSD.

## Quantifying the PDS insulation using First Passage Time Analysis

To further characterize how a PSD is isolated to ambient trafficking molecules, we explore how it can prevent random molecules to penetrate or escape the domain Ω, defined by the condensed chromatin polymer. To estimate the resident time *τ_in_* spent by Brownian molecules inside the nanoregion after crossing its boundary (fig. 3A), we run various simulations and we show this time depends weakly on the overall density of these particles or on the presence of binding sites (fig. 3B).

**FIG. 3.**
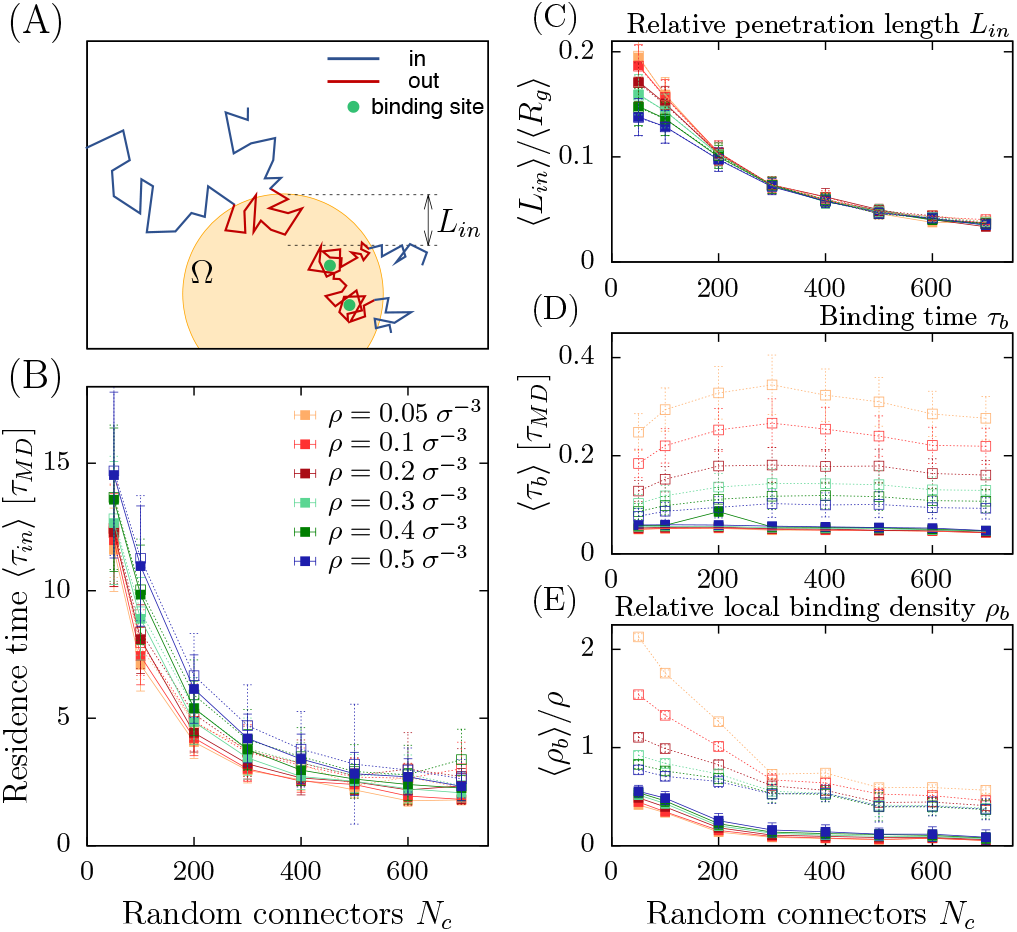
**A.** Schematic representation of molecular trajectories penetrating the phase separeted region Ω over a characteristic length *L_in_* with and without binding sites (green). **B.** Mean time 〈*τ_in_*〉 spent by a molecule inside the PSD versus number of connectors N_c_ for various densities *ρ*. Full (resp. empty) symbols indicate cases with N_bs_ = 0 (N_bs_ = 10). **C.** Ratio of the penetration length 〈*L_in_*〉 to the gyration radius 〈*R_g_*〉 versus *N_c_*. **D.** Mean binding time *τ_b_*〉 vs *N_c_*. **E.** Ratio of the local density *ρ_b_* estimated around the binding sites to the overall density *ρ* (no binding sites).

To further explore the ability of the PSD to prevent molecules from penetrating deeply inside, we defined and then estimated the penetration length *L_in_* of a trajectory before as the maximum length it can go inside the PSD before returning back to the boundary *∂*Ω. We find (fig. 3B) that on average particles cannot penetrate more than 15-20% inside even with few connectors. Furthermore, the penetration length L_in_ decays uniformly with *N_c_*.

To investigate the effects of binding sites on the retention time inside Ω, we computed the average binding time *τ_b_* of the Brownian molecules inside the region Ω and found that this time is slightly affected by the number of random connectors (fig. 3C). This result suggests an enhanced turnover of bounded particles which depends on the overall density. Finally, random connectors are sufficient to compact the polymer, leading to a partial shield of the binding sites, thus reducing the number of multiple bonds, as revealed by the local density *ρ_b_* of Brownian particles around the binding sites (fig. 3D).

## The mean escape time quantifies the insulation of PSD

Although PSDs can be isolated from the rest of their local environment, few trajectories could still escape or enter. To investigate their statistical properties, we study how single diffusing molecules positioned at the center of mass *CM* can escape. We run simulations to estimate the mean escape time 〈*τ_e_*〉 (fig. 4B) and we found a scaling law 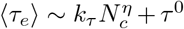, with *η* = 3.6±0.3, *k_τ_* =4 · 10^-8^ ± 10^-8^ *τ_MD_*, *τ*^0^ =35 ± 2 *τ_MD_* (no binding) and *η* = 4.0 ± 0.3, *k_τ_* = 2 · 10^-10^ ± 10^-10^ *τ_MD_*, *τ*^0^ = 80 ± 8 *τ_MD_* (with binding). The mean time *τ*^0^ is associated with the diffusing particles escaping the PSD in the absence of connectors (fig. 4A-B).

**FIG. 4.**
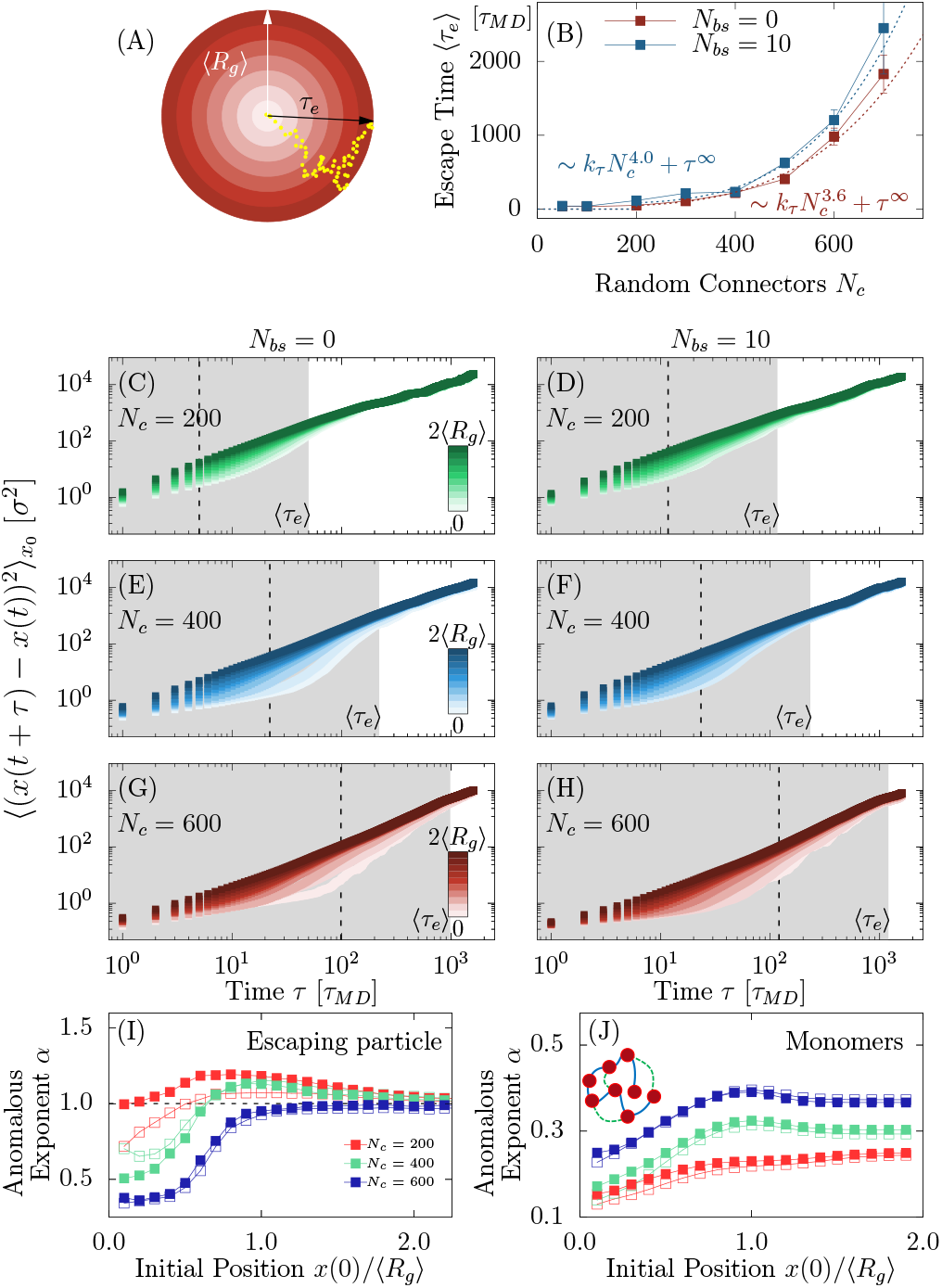
**A.** Schematic representation of a trajectory (yellow) inside the PSD, with respect to the polymer center of mass CM (color shadows). A molecule spends a random time τ_e_ before crossing the boundary. **B.** Average escaping time 〈*τ_e_*〉 from the PSD versus *N_c_* with and without binding sites. **C-H.** MSD of molecules escaping from the PSD for different values *N_c_* = 200, 400, 600, with *N_bs_* = 0 (left column) and *N_bs_* = 10 (right). Curves are colored according the range of the initial position (white inside, dark outside the PSD). Gray regions indicate the mean escape time 〈*τ_e_*〉 timescale. The binning length is 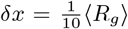. **I.** Anomalous *α*-exponent computed from the MSD of escaping particles in the time interval *τ* ∈ [1, 10^-1^*τ_e_*] with respect to the initial radial position *r*. Full (reps. empty) points correspond to *N_bs_* = 0 (resp. *N_bs_* = 10). **J.** Anomalous α-exponent computed from the MSD of monomers in the polymer center of mass reference, in the time interval *τ* ∈ [1, 10^-1^*τ_e_*] with respect to the initial radial position *r*.

To study the impact of the chromatin on diffusing particles, we also analyzed trajectories for various distances |*x*_0_| = *r* (see fig. S2) from the polymer CM and computed the average mean square displacement (MSD):

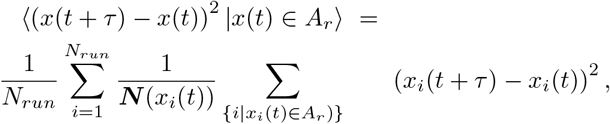

where *i* is the index of a trajectory, *x*(*t*) is the position of the trajectory inside the annulus *A_r_* = (*r, r* + *δr*) and the conditional average 〈.|*x*(*t*) ∈ *A_r_*〉 is obtained from all initial positions starting in *A_r_* at time *t*. We performed *N_run_* = 100 simulations repeated for *N_r_* = 100 polymer realizations for 2 · 10^3^ *τ_MD_* · By increasing the random connectors, a diffusing molecule trapped inside the PSD remains blocked due the many polymer loops that occupy the available space. An escape route for the diffusing particle (fig. 4B) can however emerge as a rare event, where polymer loops create a transient opening. To characterize how the polymer organization creating long-range interactions can affect the dynamics of Brownian particles, we computed the MSD functions (fig. 4C-H), showing a continuous spectrum that depends on the distance *r* from the CM and the number of connectors *N_c_*. The MSD of trajectories starting near CM (brighter curve in fig. 4C-H) shows multiple dynamics, compared to the one starting outside (darker colors). Fitting the MSD curves with ~ *D*_∞_*τ^α^*, we computed the anomalous exponents *α* for escaping molecules and also for monomers where the reference is CM. We find similar behaviors characterized by two regimes: (i) anomalous diffusion where the escaping molecules are progressively squeezed out by the polymer and (ii) normal diffusion when approaching the boundary of the PSD (see comparison in fig. 4I-J). In addition, the escape from MSD is not driven by any drift as shown in fig. S3.

## Scaling law associated with the mean escape time from a PSD

To investigate how the mean escape time for a stochastic molecule depends on the number of connectors, we use the narrow escape theory [45] allowing us to replace the moving RCL-chain that generates transient obstacle barriers by a partial reflecting boundary at the escape windows. Indeed, as suggested by the escape time results of Fig. 4B, only a small fraction of the boundary is accessible for escape. For a Brownian particle that has to escape through *N_w_* partially absorbing windows of size *a* located on a spherical surface, the escape time 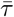 is given by [46]

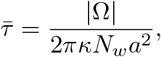

where |Ω| is the volume of the diffusing region, *κ* is partially absorbing constant that reflects the effect of the polymer on the dynamics of the moving particle. In the PSD, the accessible region Ω is the space occupied by the polymer.

Using the previous scaling laws (Fig. 1B), we aim now at estimating how the number of escaping windows *N_w_* depends on the random connectors *N_c_*. We start with the asymptotic behavior for the volume 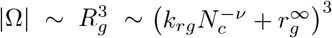, we next approximated the size of the escaping window *a* as the average pore size *ζ* (Fig. 2D), 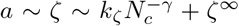. Then, we the mean escape time can be rewritten as:

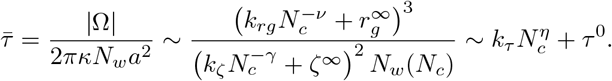

Finally we found:

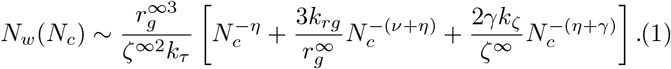

To conclude, the number of escaping windows is inversely proportional to the escaping time 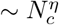.

## Concluding remarks

We demonstrated here that the PSD can result from multiple connectors that would condense chromatin fiber (Fig. 5). Using polymer model, scaling laws and numerical simulations, we found that a PSD can isolate diffusing molecules. Using a monomer resolution of 3kbp, corresponding to *σ* = 30nm, and *τ_MD_* ≈ 0.02s (used in semi-dilute polymer solutions [37]), the presence of *N_c_* ~ 50 leads to a PSD region of size 〈*R_g_*〉 ≃ 1.5 *μm*. In this context the resident time of a random particle is 〈*τ_in_*〉 ≃ 0.3s, while the escape time from the center of PSD is 〈*τ_e_*〉 ≃ 0.7s. Interestingly, these time scales are quite different from the life time of this PSD which depends on the dynamics of cross-linkers. We also reported here a boundary layer of 10-15% of the PSD size that can prevent stochastic particles from fully penetrating. Finally, we propose to use the mean escape time to quantify the ability of the PSD to retain particles inside and to measure the degree of isolation. It would be interesting to estimate the PSD organization and the mean number of cross-linkers from the distribution of anomalous exponent, extracted from future SPT experiments. This reverse engineering problem can be adressed using the *alpha*-exponent curves from figs 4I-J. Finally, the present model of beads connected by spring could be generalized in a network of interacting scaffolding proteins present in neuronal synapses at the post-synaptic density [15–17]. The ensemble produces a phase separation domain that can regulate membrane receptors. To conclude two and three dimensional polymer networks provide a mechanistic representation of phase separation that regulate local processes such as protein trafficking, transcription, plasticity and possibly many more.

**FIG. 5.**
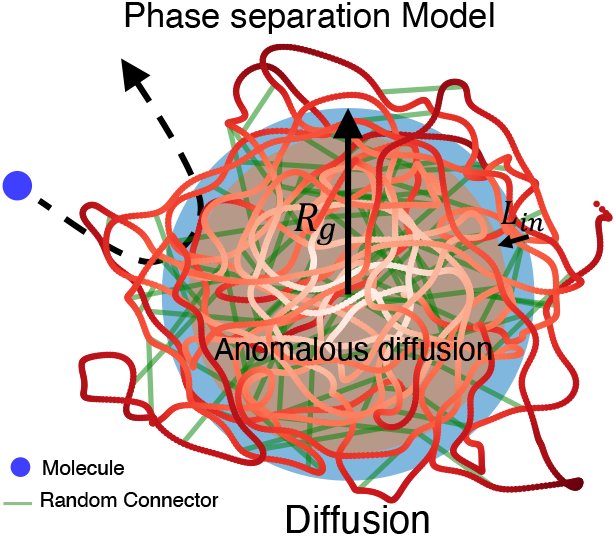
Summary of a Phase Separated Domain, described by a RCL-volume extrusion model (polymer in red), with crosslinkers (green) with a gyration radius (*R_g_*) revealing a transition area where the motion changes from anomalous diffusion to normal diffusion near the boundary (blue), characterized by the penetration length *L_in_*. A trajectory (black dashed) bounces back after penetrating of a length *L_in_*.

## Acknowledgments

A.P. is supported by a postdoctoral fellowship from the Fondation pour la Recherche Médicale (Postdoctorat en France - SPF201909009284). This project has received funding from the European Research Council (ERC) to D.H. under the European Union’s Horizon 2020 research and innovation programme (grant agreement No 882673). The authors acknowledge computational resources from Bio-Clust (IBENS) facilities.

