## Supplementary material for "Chromatin phase separated nanoregions regulated by cross-linkers and explored by single particle trajectories": so


A, Papale[1], D. Holcman[1,2]

[1] *Group of Computational Biology and Applied Mathemathics, Ecole Normale Supérieure, IBENS, Université PSL, 75005 Paris, France and*

[2] *Churchill College, University of Cambridge, CB30DS, United Kingdom*


(Dated: January 28, 2023)

The SI is separated into several sections: we first present the characteristics of the cross-linked polymer models and the associated energy. Second, we summarize our simulation procedure. Third, we expand the computation associated to the scaling law for the mean escape time from a PSD. Finally, we show that a phase separated domain does not share the characteristic of a potential well as a mechanism to retain diffusing molecules.

### GENERALIZED RANDOM CROSS-LINKER POLYMER MODEL TO DESCRIBE DENSE CHROMATIN PHASES

#### Construction of the polymer chain from potential well

We present here an extension of the random cross-linker model [1] that includes volume excluded interactions. This extension uses bead-spring polymer model, originating from the Kremer-Grest [2] coarse-grained model [3–5]. The model is constructed as follows: we consider a bead-spring polymer with a total of $N_{mon}$ monomers where we have added $N_c$ cross-linkers located at random positions. Each of $N_{mon}$ interacting monomer of the polymer chain corresponds to 3 kbp, with size $\sigma = 30nm$ and their dynamics is described by the potential energy which is the sum of several terms for the vector position of all beads $(\vec{r}_1, ..\vec{r}_N)$:

1. **The Lennard-Jones potential** $U_{LJ}(\vec{r}_1, ..\vec{r}_N)$ describes the excluded volume interactions. We took for $U_{LJ}$ a truncated and shifted Lennard-Jones potential: two beads repel when their distance is less than $2^{1/6}\sigma$, which corresponds to the minimum of the potential:

$$U_{LG}(r) = \begin{cases} 4\epsilon \left[ \left(\frac{\sigma}{r}\right)^{12} - \left(\frac{\sigma}{r}\right)^{6} + \frac{1}{4} \right] & r \leq r_c \\ 0 & r > r_c, \end{cases} \tag{S1}$$

where $r$ is the distance between any two monomers while the cutoff distance $r_c = 2^{1/6}\sigma$ conserves only the repulsive contribution. The energy scale is $\epsilon = \kappa_B T$, where $T = 300$ K.

2. **Non-linear elastic potential (FENE).** The linear connectivity of the chain is ensured by bonding nearest-neighbours monomers with the finitely extensible non-linear elastic potential (FENE): the energy $U_{FENE}(\vec{r}_1, ..\vec{r}_N)$ is associated to the backbone of the polymer chain. This potential enforces the connectivity of the chain, so that two consecutive particles cannot be distant by more than $R_0 = 1.5\sigma$.

$$U_{FENE}(r) = \begin{cases} -0.5\kappa R_0^2 \ln\left(1 - (r/R_0)^2\right) & r \leq r_c \\ \infty & r > r_c, \end{cases} \tag{S2}$$

where $\kappa = 30\epsilon/\sigma^2$ is the spring constant and $R_0 = 1.5\sigma$ is the maximum extension of the elastic FENE bond.

3. **The bending energy $U_{bend}$.** The stiffness of the polymer is quantified by the bending energy which depends on the cosine of the angle between two consecutive bonds along the chain. The bending energy $U_{bend}(\vec{r}_1, ..\vec{r}_N)$ penalizes consecutive bond vectors $\vec{b}_i = \vec{r}_{i+1} - \vec{r}_i$ that are not parallel. Using the monomers positions $\vec{r}_i$ along the chain, the analytical expression is given by

$$U_{\text{bend}}(\vec{r}_{i-1}, \vec{r}_i, \vec{r}_{i+1}) = \kappa_{\text{bend}} \left( 1 - \frac{(\vec{r}_{i+1} - \vec{r}_i) \cdot (\vec{r}_i - \vec{r}_{i-1})}{|\vec{r}_{i+1} - \vec{r}_i| \, |\vec{r}_i - \vec{r}_{i-1}|} \right), \tag{S3}$$

where $\kappa_\theta = 5\kappa_B T$ is the bending constant as the Kuhn's length of the 30-nm fiber is $l_K = 300$ nm.

4. **Harmonic potential $U_{harm}$ between random connectors.** The presence of loops is implemented with an harmonic potential to add $N_c$ cross-linkers between randomly chosen monomers. The energy is given by

$$U_{harm}(r_{i,j}) = \frac{k_{rc}}{2} r_{i,j}^2, \quad \text{(S4)}$$

where $k_{rc} = 0.5\sigma^2/\epsilon$ is the spring constant, $r_{i,j} = |\vec{r}_i - \vec{r}_j|$ the distance between two non-nearest-neighbours monomers connected by a random connector.

To summarize the polymer chain is described by the following energy term:

$$H_{INT}(r) = \sum_{i,j}^{N_{mon}} U_{LG}(\vec{r}_1, ..\vec{r}_N) + \sum_{i=1}^{N_{mon}-1} U_{FENE}(r_{i,i+1}) + \sum_{i=2}^{N_{mon}-1} U_{\text{bend}}(\vec{r}_{i-1}, \vec{r}_i, \vec{r}_{i+1}) + \sum_{k=(k_i,k_j)}^{N_c} U_{harm}(r_{k_i,k_j}). \quad \text{(S5)}$$

#### Langevin's dynamics of the polymer chain

The dynamics of the chain is described by the Langevin equation:

$$m\frac{dv}{dt} = -m\gamma v - \nabla H_{INT} + \sqrt{2dD}\dot{\eta}. \quad \text{(S6)}$$

where $\eta$ is zero-mean Gaussian noise. We recall that $N_{mol}$ molecules and $N_{mon}$ monomers of size $\sigma$ diffuse with diffusion coefficient $D = \frac{\kappa_B T}{\gamma}$. The molecule-molecule and molecule-monomer interactions are defined according the truncated Lennard-Jones potential S1. Along the chain, we positioned $N_{bs}$ binding sites on monomers: a free molecule can then be attached to a binding site when their relative distance is $d < 2 \cdot 2^{1/6}\sigma$ via a Lennard-Jones attractive potential with $\epsilon = 5\kappa_B T$. A molecule can attach to only one binding site, while each binding site can accommodate more than one binding molecule.

### NUMERICAL IMPLEMENTATION

The model has been investigated performing fixed-volume and constant-temperature Molecular Dynamics (MD) simulations with implicit solvent. The equations of motion are integrated using a velocity Verlet algorithm and Langevin thermostat with temperature $T = \kappa_B$ and damping constant $\gamma = 0.5\tau_{MD}^{-1}$ where $\tau_{MD} = \sigma(m/\epsilon)^{1/2}$ is the Lennard-Jones time scale. In the case of semi-dilute polymer solutions, it is equivalent to $\tau_{MD} \approx 0.02$ s [3].
The integration time step is set to $\Delta t = 5 \cdot 10^{-3}\tau_{MD}$. The length of each MD run for the system composed by an already equilibrated RCL-polymer and the diffusive particles is equal to $5 \cdot 10^6$ simulation steps ($2.5 \cdot 10^4 \tau_{MD}$) after an equilibrium run of $10^6$ simulation steps. The effect of random cross-linking is obtained by considering $10^2$ different random polymer connectivities.

### SCALING LAW FOR THE MEAN ESCAPE TIME FROM A PSD

The escape time $\bar{\tau}$ for a Brownian particle escaping through $N_w$ partially absorbing windows of size $a$ located on a spherical surface, is given by [6]:

$$\bar{\tau} = \frac{|\Omega|}{2\pi\kappa N_w a^2},$$

where $|\Omega|$ is the volume of the diffusing region, $\kappa$ is partially absorbing constant that reflects the effect of the polymer on the dynamics of the moving particle. In the PSD, the accessible region $\Omega$ is the space occupied by the polymer. We show here the detailed computations for estimating how the number of escaping windows $N_w$ depends on the random connectors $N_c$. We start with the asymptotic behavior for the volume $|\Omega| \sim R_g^3 \sim \left(k_{rg}N_c^{-\nu} + r_g^{\infty}\right)^3$, we next approximated the size of the escaping window $a$ as the average pore size $\zeta$, $a \sim \zeta \sim k_\zeta N_c^{-\gamma} + \zeta^{\infty}$. Then, we the mean escape time can be rewritten as:

$$\bar{\tau} = \frac{|\Omega|}{2\pi\kappa N_w a^2} \sim \frac{\left(k_{rg}N_c^{-\nu} + r_g^{\infty}\right)^3}{\left(k_\zeta N_c^{-\gamma} + \zeta^{\infty}\right)^2 N_w(N_c)} \sim k_\tau N_c^{\eta} + \tau^0.$$

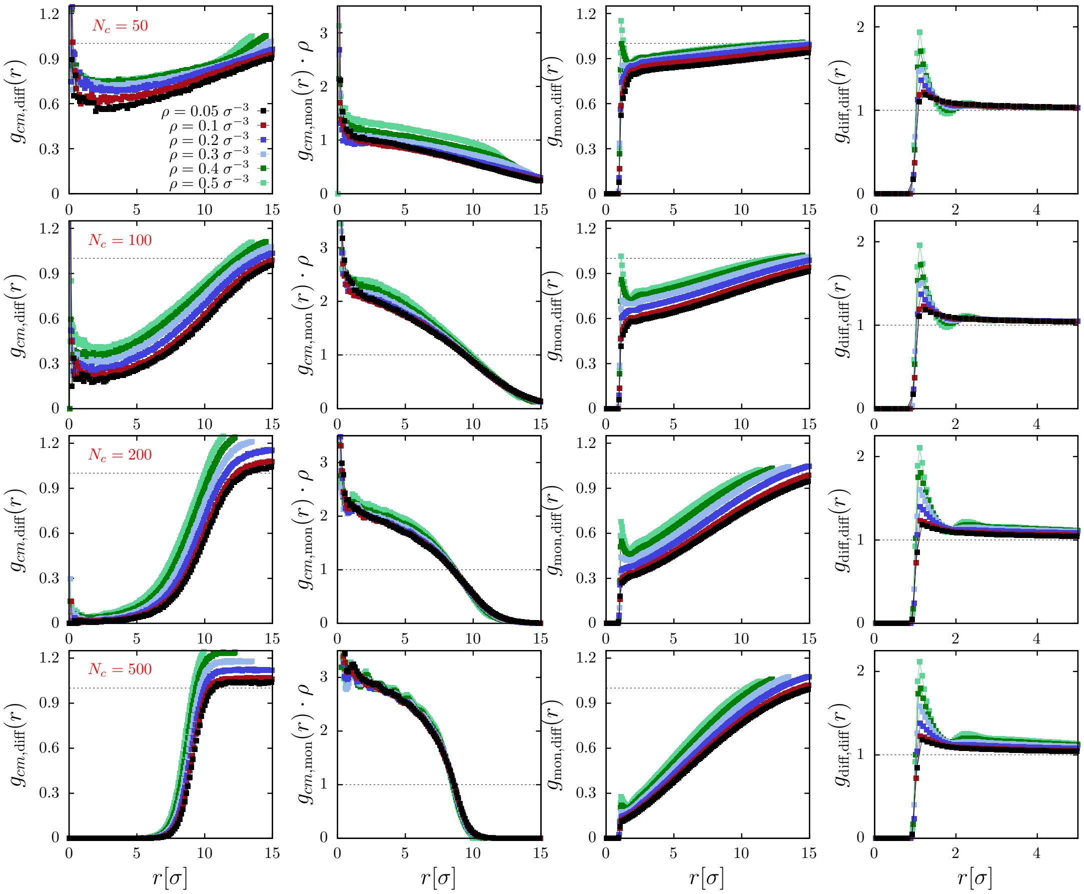


FIG. S1. First column: molecules radial distribution function $g_{\mathrm{mol}}(r)$ for different density; Second column: polymer radial distribution function $g_{\mathrm{mon}}(r)$. Third column: molecules-monomers pair correlation function $g_{\mathrm{mol}}(r)$. Fourth column: molecule-molecule pair correlation function $g_{\mathrm{mol,mol}}(r)$. Fifth column: monomer-monomer pair correlation function $g_{\mathrm{mon,mon}}(r)$.

We can the isolate the espression for the number of escaping windows as a funtion of the number of connectors $N_c$:

$$N_w(N_c) \sim \frac{k_{rg}^3 N_c^{-3\nu} + 3k_{rg}^2 N_c^{-2\nu} r_g^{\infty} + 3k_{rg} N_c^{-\nu} r_g^{\infty 2} + r_g^{\infty 3}}{\left(k_\zeta^2 N_c^{-2\gamma} + \zeta^{\infty 2} + 2k_\zeta N_c^{-\gamma}\zeta^{\infty}\right)\left(k_\tau N_c^{\eta} + \tau^{\infty}\right)}$$

that can be rearranged as

$$N_w(N_c) \sim \frac{r_g^{\infty 3}}{\zeta^{\infty 2}\tau^{\infty}} \left[\frac{1 + \frac{3k_{rg}}{r_g^{\infty}} N_c^{-\nu} + O\left(N_c^{-2\nu}\right)}{\left(1 + \frac{2k_\zeta}{\zeta^{\infty}} N_c^{-\gamma} + O\left(N_c^{-2\gamma}\right)\right)\left(1 + \frac{k_\tau}{\tau^{\infty}} N_c^{\eta}\right)}\right].$$

In the limit $N_c$ large we can set

$$1 + \frac{k_\tau}{\tau^{\infty}} N_c^{\eta} \sim \frac{k_\tau}{\tau^{\infty}} N_c^{\eta}$$

and get

$$N_w(N_c) \sim \frac{r_g^{\infty 3}}{\zeta^{\infty 2} k_\tau} \left[\frac{N_c^{-\eta} + \frac{3k_{rg}}{r_g^{\infty}} N_c^{-(\nu+\eta)} + O\left(N_c^{-2\nu}\right)}{1 + \frac{2k_\zeta}{\zeta^{\infty}} N_c^{-\gamma} + O\left(N_c^{-2\gamma}\right)}\right].$$

Expanding the denominator we then get

$$N_w(N_c) \sim \frac{r_g^{\infty 3}}{\zeta^{\infty 2} k_\tau} \left(N_c^{-\eta} + \frac{3k_{rg}}{r_g^{\infty}} N_c^{-(\nu+\eta)} + O\left(N_c^{-2\nu}\right)\right) \left[1 + \left(\frac{2\gamma k_\zeta}{\zeta^{\infty}} N_c^{-\gamma} + O\left(N_c^{-2\gamma}\right)\right)\right].$$

Finally, we get

$$N_w(N_c) \sim \frac{r_g^{\infty 3}}{\zeta^{\infty 2} k_\tau} \left( N_c^{-\eta} + \frac{3k_{rg}}{r_g^{\infty}} N_c^{-(\nu+\eta)} + \frac{2\gamma k_\zeta}{\zeta^{\infty}} N_c^{-(\eta+\gamma)} \right).$$

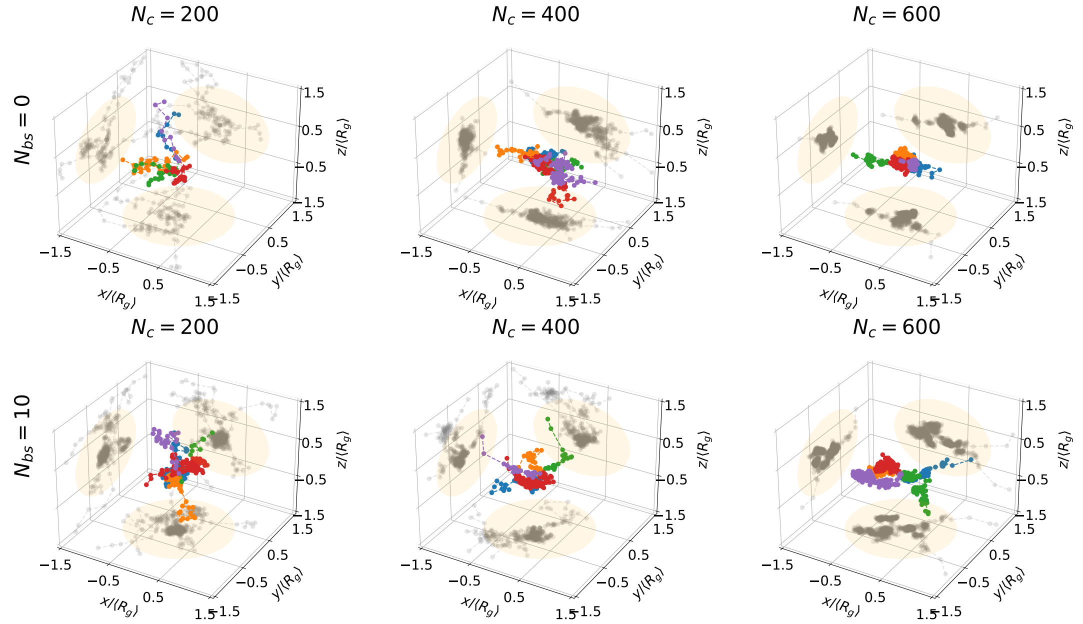


FIG. S2. Few examples of trajectories of escaping particles for systems with $N_c = 200, 400, 600$ (columns) and $N_{bs} = 0, 10$ (rows) highlighted with different colors. On each surface the projected trajectories are shown in gray, orange circles represent the projections of the $\Omega$ regions defined by the gyration radius.

### MECHANISM TO RETAIN DIFFUSING MOLECULES IS NOT A POTENTIAL WELL

#### Main definitions for recovering a potential well from trajectories

To investigate whether the PSD can retain stochastic particles with the characteristic of a potential well, we assumed that trajectories could result from a coarser spatio-temporal motion following the stochastic process [7, 8]

$$\dot{X} = a(\boldsymbol{X}) + \sqrt{2}B(\boldsymbol{X})\dot{W}, \tag{S7}$$

where $a(\boldsymbol{X})$ is the drift field and $B(\boldsymbol{X})$ is a matrix and $\dot{W}$ is a random noise. The drift in eq. S7 can be recovered from SPTs acquired at any infinitesimal time step $\Delta t$ by estimating the conditional moments of the trajectory displacements $\Delta\boldsymbol{X} = \boldsymbol{X}(t + \Delta t) - \boldsymbol{X}(t)$ [8–11]

$$a(x) \;=\; \lim_{\Delta t \to 0} \frac{\mathbb{E}[\Delta\boldsymbol{X}(t) \,|\, \boldsymbol{X}(t) = x]}{\Delta t}, \tag{S8}$$

$$\tag{S9}$$

The notation $\mathbb{E}[\cdot \,|\, \boldsymbol{X}(t) = x]$ represents averaging over all trajectories that are passing at point $x$ at time $t$. To estimate the local drift $a(\boldsymbol{X})$ at each point $\boldsymbol{X}$ and at a fixed time resolution $\Delta t$, we use a procedure based on a square grid. The local estimators to recover the vector field consist in grouping points of trajectories within a lattice of square bins $S(x_k, \Delta x)$ centered at $x_k$ and of width $\Delta x$. For an ensemble of $N$ three-dimensional trajectories $\{\boldsymbol{X}_i(t_j) = (x_i^{(1)}(t_j), x_i^{(2)}(t_j)), x_i^{(3)}(t_j)i = 1..N, j = 1..M_i\}$ with $M_i$ the number of points in trajectory $\boldsymbol{X}_i$

and successive points recorded with an acquisition time $t_{j+1} - t_j = \Delta t$. The discretization of eq. S8 for the drift $a(x_k) = (a^{(1)}(x_k), a^{(2)}(x_k), a^{(3)}(x_k))$ in a bin centered at position $x_k$ is

$$a^{(u)}(x_k) \approx \frac{1}{N_k} \sum_{i=1}^{N} \sum_{j=0, x_i(t_j)\in S(x_k,\Delta x)}^{M_i-1} \left( \frac{x_i^{(u)}(t_{j+1}) - x_i^{(u)}(t_j)}{\Delta t} \right), \tag{S10}$$

where $u = 1..3$ and $N_k$ is the number of points $x_i(t_j)$ falling in the square $S(x_k, r)$.
At this stage, we would like to compare the empirical drift obtained from the trajectories of diffusing particles with the one generated by a parabolic well. We consider the basin of attraction of a truncated elliptic parabola with the associated energy function

$$U(\boldsymbol{X}) = \begin{cases} A\left[\left(\frac{x^{(1)} - \boldsymbol{\mu}^{(1)}}{a}\right)^2 + \left(\frac{x^{(2)} - \boldsymbol{\mu}^{(2)})}{b}\right)^2 + \left(\frac{x^{(3)} - \boldsymbol{\mu}^{(3)})}{c}\right)^2 - 1\right], & \boldsymbol{X} \in \mathcal{B} \\ 0 & \text{otherwise} \end{cases} \tag{S11}$$

where $A > 0$ and $\boldsymbol{X} = [x^{(1)}, x^{(2)}, x^{(3)}]$, $\boldsymbol{\mu} = [\boldsymbol{\mu}^{(1)}, \boldsymbol{\mu}^{(2)}, \boldsymbol{\mu}^{(3)}]$ is the center of the well, $a, b, c$ are the elliptic semi-axes lengths and the elliptic boundary is defined by

$$\mathcal{B} = \{\boldsymbol{X}\text{such that } A\left[\left(\frac{x^{(1)} - \boldsymbol{\mu}^{(1)}}{a}\right)^2 + \left(\frac{x^{(2)} - \boldsymbol{\mu}^{(2)})}{b}\right)^2 + \left(\frac{x^{(3)} - \boldsymbol{\mu}^{(3)})}{c}\right)^2 - 1\right] = 0\}. \tag{S12}$$

The PSD is centered at $\boldsymbol{\mu}^{(1)} = \boldsymbol{\mu}^{(2)} = \boldsymbol{\mu}^{(3)} = 0$ and the elliptic semi-axes lengths are approximated by the radius gyration $R_g$. To estimate the attraction coefficient $A$, we use the least-square regression formula

$$A = R_g^2 \frac{1}{2} \frac{\sum_{k=1..3, i=1}^{M} a^{(k)}(\boldsymbol{X}_i) x_i^{(k)}}{\sum_{k=1}^{3} \sum_{i=1}^{M} (x_i^{(k)})^2}, \tag{S13}$$

where $\boldsymbol{X}_i = [x_i^{(1)}, x_i^{(2)}, x_i^{(3)}]$ $(i = 1 \dots M)$ are the centers of the $M$ bins.
Finally, we can estimate the quality of the well (parabolic index) based on the residual least square error:

$$S = 1 - \frac{1}{2} \frac{\left(\sum_{k=1..3, i=1}^{M} a^{(k)}(\boldsymbol{X}_i) x_i^{(k)}\right)^2}{\left(\sum_{k=1}^{3} \sum_{i=1}^{M} (x_i^{(k)})^2\right)\left(\sum_{i=1}^{M} ||a(\boldsymbol{X}_i)||^2\right)}. \tag{S14}$$

The index $S \in [0, 1]$ is defined such that $S \to 0$ for a drift field generated by a parabolic potential well and $S_k \to 1$ for a random drift vector field as observed for diffusive motion.

#### Application: estimation of the vector field associated to trajectories inside the PSD

We apply the previous procedure to recover and characterise a possible drift field inside the PSD. We found that there was no drift associated with the PSD, as summarized in table below. Indeed, the score parameter $S \approx 1$ for the

| $N_c$ | $N_{bs} = 0$ | $N_{bs} = 10$ |
|---|---|---|
| 200 | $A = 2 \cdot 10^{-4}$ | $A = 6 \cdot 10^{-5}$ |
| 400 | $A = 10^{-4}$ | $A = 9 \cdot 10^{-5}$ |
| 600 | $A = 5 \cdot 10^{-5}$ | $A = 10^{-4}$ |

TABLE S1. A values computed from the simulated trajectories described in fig. S3

different parameter values reported in the table. These results show that the PSD (Fig. S3) traps stochastic particles with a mechanism different from an attracting potential well.

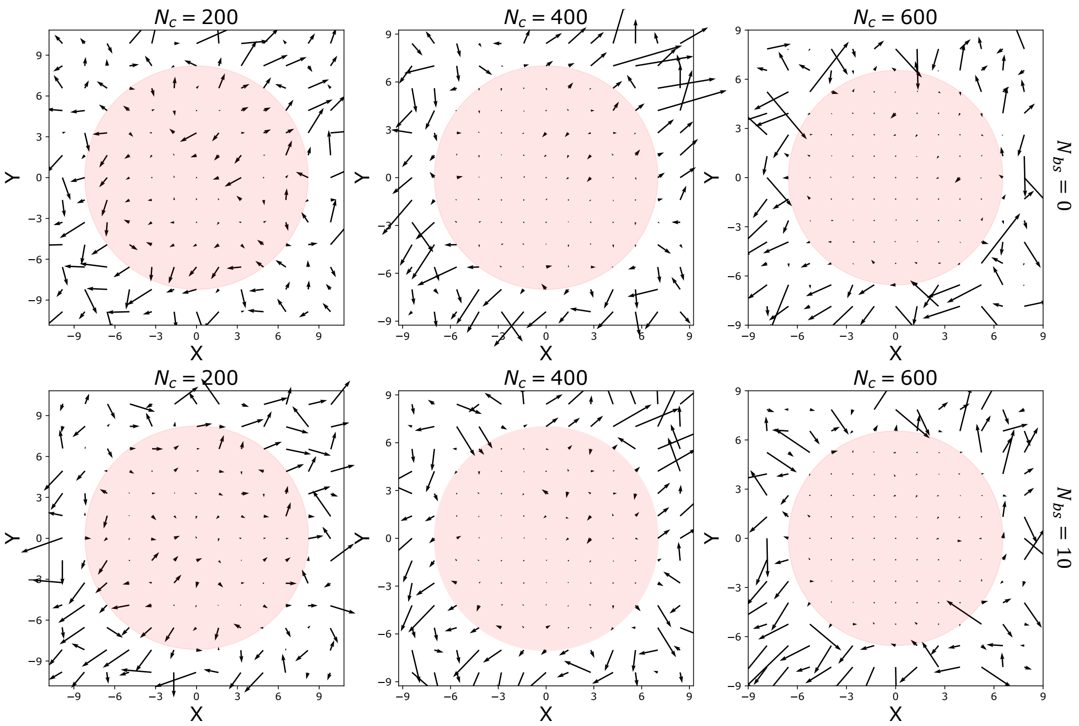


FIG. S3. Vector fields on the plane $x-y$, $z=0$, computed from escaping particle trajectories for systems with $N_c = 200, 400, 600$ (columns) and $N_{bs} = 0, 10$ (rows). Circles represent the projections of the $\Omega$ regions defined by the gyration radius.

[11] N. Hozé and D. Holcman, "Statistical methods for large ensembles of super-resolution stochastic single particle trajectories in cell biology," 2017.
